# Assigning mutational signatures to individual samples and individual somatic mutations with SigProfilerAssignment

**DOI:** 10.1101/2023.07.10.548264

**Authors:** Marcos Díaz-Gay, Raviteja Vangara, Mark Barnes, Xi Wang, S M Ashiqul Islam, Ian Vermes, Nithish Bharadhwaj Narasimman, Ting Yang, Zichen Jiang, Sarah Moody, Sergey Senkin, Paul Brennan, Michael R Stratton, Ludmil B Alexandrov

## Abstract

Analysis of mutational signatures is a powerful approach for understanding the mutagenic processes that have shaped the evolution of a cancer genome. Here we present SigProfilerAssignment, a desktop and an online computational framework for assigning all types of mutational signatures to individual samples. SigProfilerAssignment is the first tool that allows both analysis of copy-number signatures and probabilistic assignment of signatures to individual somatic mutations. As its computational engine, the tool uses a custom implementation of the forward stagewise algorithm for sparse regression and nonnegative least squares for numerical optimization. Analysis of 2,700 synthetic cancer genomes with and without noise demonstrates that SigProfilerAssignment outperforms four commonly used approaches for assigning mutational signatures. SigProfilerAssignment is freely available at https://github.com/AlexandrovLab/SigProfilerAssignment with a web implementation at https://cancer.sanger.ac.uk/signatures/assignment/.

## MAIN

Somatic mutations accumulate in the genomes of all cells of the human body throughout an individual’s lifetime^1,2^. These mutations arise from different endogenous and exogenous mutational processes, with each process generating a characteristic pattern of mutations, known as a *mutational signature*^3^. By leveraging the vast amounts of high-throughput DNA sequencing data generated over the last two decades, distinct mutational signatures have been elucidated from various cancer types^4,5^ and normal somatic tissues^6-9^. Sets of mutation-type specific reference signatures have been developed and deposited in the Catalogue of Somatic Mutations in Cancer (COSMIC) database^4,10^, including signatures of single base substitutions (SBSs), doublet base substitutions (DBSs), small insertions and deletions (IDs), and copy number alterations (CNs). In practice, mutational signatures have been widely utilized in research and clinical settings, including for detecting previously unappreciated cancer predispositions^11,12^, pathogenic classification of germline variants^13^, clinical management of cancer patients^14^, and identifying sensitivity to anti-cancer therapeutics^15^.

There are at least two distinct approaches for analyzing mutational signatures. *De novo* extraction is an unsupervised machine learning approach that allows identifying the patterns of known and previously unknown mutational signatures^16^. This type of analysis is predominately used for deriving reference signatures as it requires large cohorts, generally more than 100 samples and, usually, many thousands of samples. In contrast, *refitting* of mutational signatures is a numerical optimization approach that allows the assignment of known (in most cases, reference) signatures to an individual sample by quantifying the number of mutations attributed to each signature operative in that sample. While refitting cannot identify or quantify the activities of previously unknown mutational signatures, this approach is widely applied to small cohorts and for clinical samples where evaluations are almost exclusively performed for an individual cancer patient^17^.

In the past decade, multiple tools for refitting known mutational signatures were developed, including deconstructSigs^17^, MutationalPatterns^18,19^, sigLASSO^20^, and SignatureToolsLib^5,21^. The majority of these tools lack online interface and provide support almost exclusively for substitution signatures. Further, most tools for refitting of known mutational signatures have never been compared and no existing tool supports probabilistically assigning mutational signatures to somatic mutations. To address these limitations, here, we present SigProfilerAssignment, a comprehensive computational tool for assigning mutational signatures to individual samples and individual somatic mutations (**Fig. 1**). In contrast to other tools, SigProfilerAssignment provides desktop and online support for all types of mutational signatures, including the COSMIC sets of reference SBS, DBS, ID, and CN signatures. In addition to COSMIC reference signatures, SigProfilerAssignment supports assignment of *de novo* extracted mutational signatures as well as of a user provided set of custom mutational signatures. Our benchmarking based on 2,700 simulated cancer genomes demonstrates that SigProfilerAssignment outperforms other commonly used tools on simulation data with and without noise.

**Figure 1.**
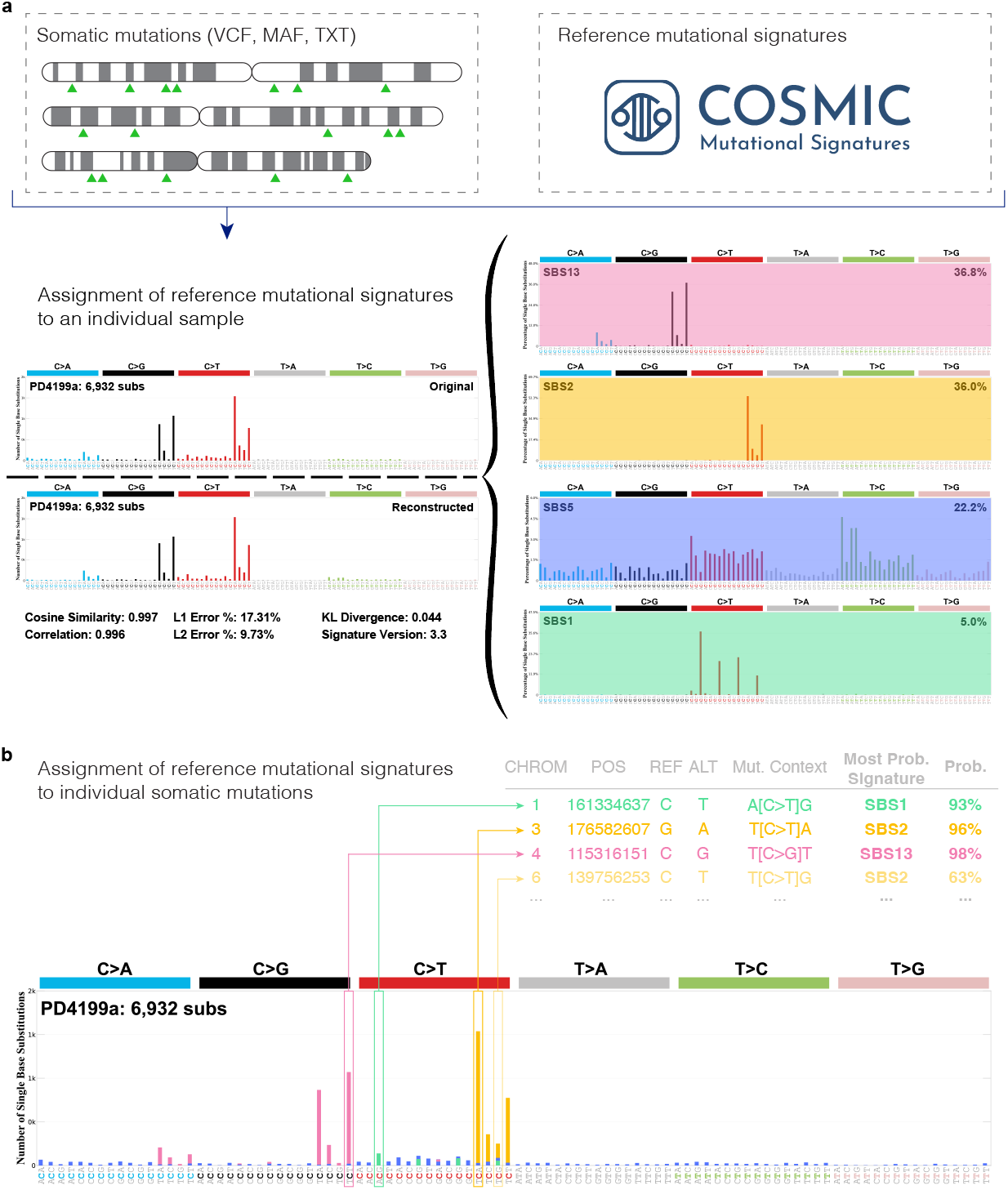
Assigning known mutational signatures to an individual sample and individual mutations with SigProfilerAssignment. SigProfilerAssignment supports input data in a standard format (VCF, MAF, or text) and it allows assigning a set of known signatures (*e*.*g*., ones from the COSMIC database) to an ***a)*** individual sample and ***b)*** probabilistically to an individual somatic mutation. Note that the probabilistic assignment of mutational signatures to an individual somatic mutation is only possible if a user provides a list of individual mutations (*e*.*g*., VCF file) for the examined sample instead of a mutational vector, as a mutational vector lacks information for individual mutations.

Given a set of known mutational signatures and a set of mutations in a cancer genome, both classified under the same mutational schema^3,22^, SigProfilerAssignment identifies the number of mutations caused by each signature in that cancer genome (**Fig. 1*a***). Mathematically, a mutational schema can be represented as a finite alphabet Ξ of mutation types containing a total of ξ letters. Here, a mutational signature is defined as a probability mass function with domain the alphabet Ξ.

In vector notation, a mutational signature can be denoted as 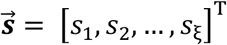, where *s*_*k*_, 1 ≤*k* ≤ *ξ*, is the probability for the mutational signature,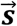, to cause mutations of type corresponding to the *k*^*th*^ letter of the alphabet Ξ. Since a mutational signature is a probability mass function, 0 ≤ s_*k*_ ≤ 1 and 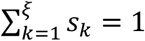. As such, a set of known *n* mutational signatures can be expressed as a signature matrix, 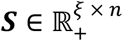, where 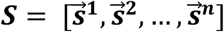. Further, a set of mutations in a cancer genome can be defined as 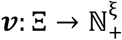. In vector notation, a set of mutations in a cancer genome 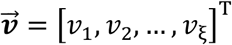, where *ξ*_*k*_, 1 ≤ *k* ≤ *ξ*, reflects the number of mutations in that cancer genome of the mutation type corresponding to the *k*^*th*^ letter of the alphabet Ξ. SigProfilerAssignment takes as an input a signature matrix, ***s***, and a set of mutations, 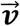, to output a column vector of activities 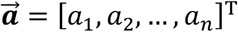, where *a*_*t*_ 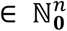, 1 ≤ *t* ≤ *n*, corresponding to the number of somatic mutations attributed to the *t*^th^ mutational signature. The underlying assumption of assigning mutational signatures is that the mutations within a sample can be approximated as a superposition of known mutational signatures and their activities:

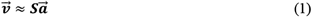

Thus, subject to 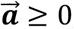, one needs to derive the vector 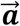 that best fits the provided input data. To solve this optimization problem, SigProfilerAssignment uses a custom implementation of the forward stagewise algorithm^23^ and it applies nonnegative least squares (NNLS)^24^, based on the Lawson-Hanson method^24^:

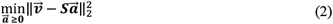

The algorithm starts by first computing a minimum relative error, 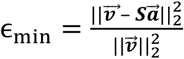, by deriving the optimal nonnegative vector 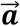 for the complete set of all reference signatures, ***S***, using equation *(2)*. This minimum error provides the best possible explanation of the data, but it also results in overfitting as all available signatures are utilized. Next, the tool uses steps for removing and adding signatures based on the backward and forward stepwise algorithms, respectively^23^. First, signatures are removed by employing a backward stepwise algorithm^23^ (**Algorithm 1**). Specifically, each signature from the reference signature set, ***S***, is removed iteratively and the remaining signature set, 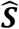, is attributed to the sample 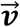 by applying equation *(2)*. The increase in the relative error, 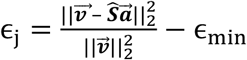 due to removing a signature is calculated by adding the *j*^th^ signature from ***S***. The signature with the least relative increase in error rate is removed from the signature set, ***S*** provided that the increase is less than a specific threshold (default value of 0.01). After the final removal of the signature with least relative error rate increase, the minimum relative error, ϵ_min_, and the set of signatures, ***S***, are updated to reflect this removal. The removal step is repeated until all signatures satisfying the conditions are removed from ***S***. The removal steps are followed by addition steps based on the forward stepwise algorithm23 (Algorithm 1). Specifically, each of the previously removed reference signatures is added back iteratively to .. and the new signature set, 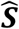, is fit for the sample 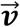 by applying equation (2). Thus, the decrease in the relative error, 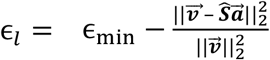, due to adding a signature is calculated by adding the *l*^th^ signature to ***S***. The signature with maximum relative decrease of the error rate is added back to the signature set, ***S***, provided that the increase is more than a specific threshold (default value of 0.05). After the final addition of the signature with most relative rate decrease, the minimum relative error, ϵ_678_, and the set of signatures, ***S***, are updated to reflect this addition. The addition step is repeated until all signatures satisfying the conditions are added back to ***s***. Lastly, the addition and removal steps are repeated until convergence, where no signature is added or removed from the list of signatures (**Algorithm 1**).

In addition to quantifying the activity of each mutational signature, SigProfilerAssignment also assigns known signatures to individual mutations (**Fig. 1*b***) based on their specific mutational context:

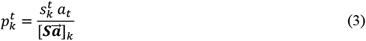

where, 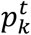 represents the probability of a mutation corresponding to the *k*^*th*^ letter of the alphabet Ξ being caused by the *t*^*th*^ signature in the sample; 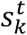 is the probability of the *t*^*th*^ signature to cause mutation corresponding to the *k*^*th*^ letter of the alphabet Ξ; *a*_*t*_ is the number of mutations attributed to the *t*^*th*^ mutational signature; and 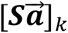 is the value of the *k*^*th*^ element of the vector obtained by the matrix multiplication of the signature matrix, ***s***, and the derived signature activities, 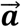.

To evaluate the performance of SigProfilerAssignment and another four commonly used tools for refitting mutational signatures^5,17-21^, we performed a comparative benchmarking using a previously generated independent synthetic dataset^16^ (**Fig. 2**). The dataset encompasses the SBS patterns of 2,700 simulated cancer genomes, corresponding to 300 tumors from nine different cancer types, generated using 21 different COSMIC reference signatures. To emulate a typical refitting of mutational signatures, the complete set of 79 COSMICv3.3 SBS signatures was used as input. The mutational signature activities obtained by each tool were compared against the ground truth activities used to synthetically generate these samples. Three different levels of random noise (0%, 5%, and 10%) were tested to assess the stability of the different algorithms in a real biological context. To evaluate the accuracy of the signature refitting, we calculated sensitivity, specificity, and F_1_ score (**Methods**). In addition, we also examined the runtime and memory utilization of each tool.

**Figure 2.**
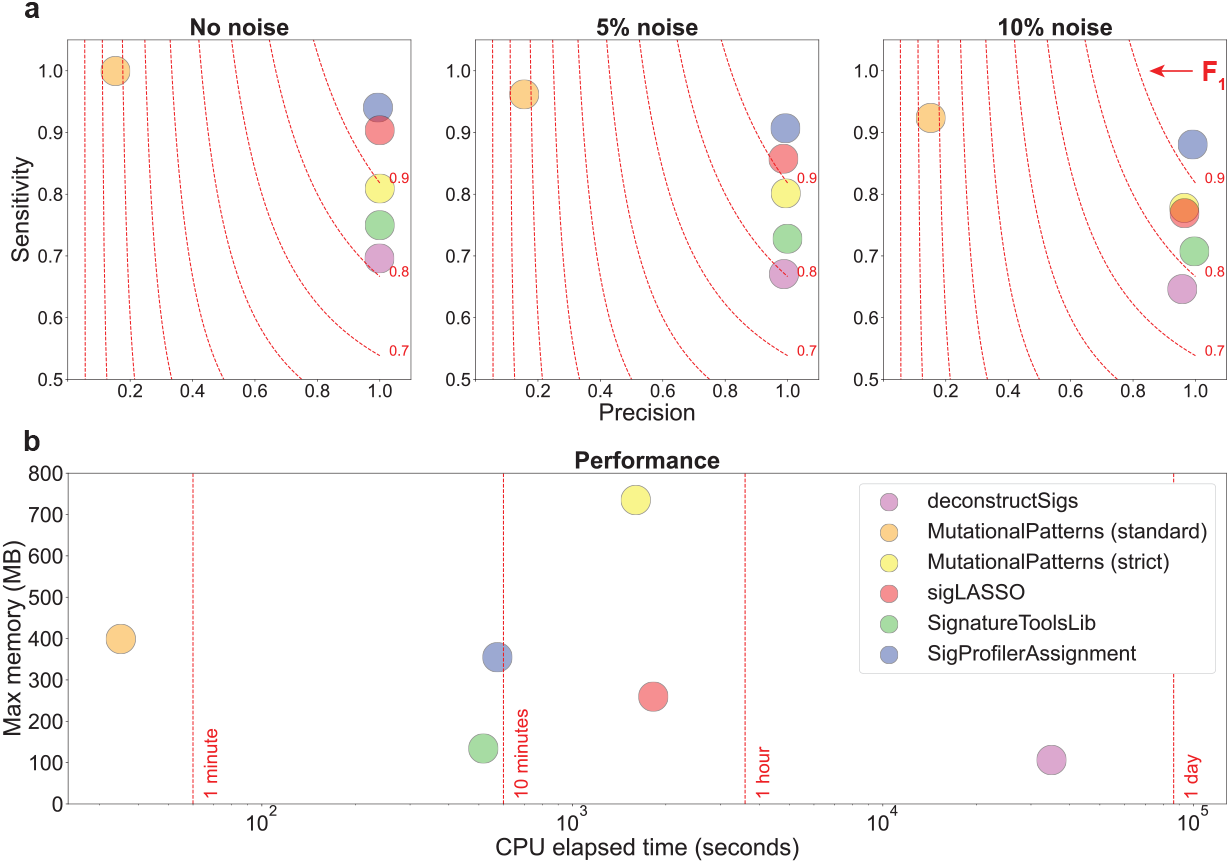
Benchmarking of SigProfilerAssignment and four other tools for assigning mutational signatures. Each tool was evaluated using 2,700 synthetic cancer genomes generated using 21 different COSMIC reference mutational signatures. All COSMICv3.3 signatures were used as the input set of known mutational signatures. ***a)*** Three different levels of non-systematic random noise (0%, 5%, and 10%) were used to evaluate the precision (x-axes), sensitivity (y-axes), and F_1_ scores (harmonic mean of precision and sensitivity; red dotted lines) of each tool. ***b)*** Computational benchmarking based on CPU elapsed time (x-axis; log-scaled) and maximum memory usage (y-axis) for each tool.

Our synthetic benchmarking revealed that SigProfilerAssignment outperforms all other approaches for the examined noise levels (**Fig. 2*a***). For 10% random noise, only SigProfilerAssignment obtained an F_1_ score >0.90. In all cases, SigProfilerAssignment exhibited a high precision while showing an improved sensitivity compared to other approaches (**Fig. 2*a***). In terms of computational performance, SigProfilerAssignment processed the 2,700 samples within 9.6 minutes (0.21 seconds per sample). Only the standard mode of MutationalPatterns generated results substantially faster (**Fig. 2*b***). However, MutationalPatterns’ standard mode exhibited sub-optimal performance, with a significant drop in precision for all noise levels, likely due to overfitting of the input data (**Fig 2*a***)^19^. This issue has been addressed in the most recent version of MutationalPatterns with the addition of a strict mode^18^, albeit with a significant computational performance cost (**Fig. 2*b***). Other approaches limit overfitting by implementing different penalties based on the L1 error (*viz*., sigLASSO)^20^ or the sum-squared error (*viz*., deconstructSigs)^17^, as well as post-hoc filters based on the percentage of the total number of mutations attributed to a given signature (*viz*., deconstructSigs and SignatureToolsLib)^5,17^. No significant memory requirements were observed for any of the tools (**Fig. 2*b***).

Assigning mutational signatures to individual samples provides an opportunity to identify the processes responsible for somatic mutations on a sample-by-sample basis. Considering our synthetic benchmarking, SigProfilerAssignment stands out as the most precise and sensitive tool while maintaining high computational performance and bringing novel capabilities. To the best of our knowledge, SigProfilerAssignment represents the first computational tool for assigning signature probabilities to individual mutations, which can allow uncovering the mutational processes responsible for specific driver genomic alterations leading to tumor evolution. SigProfilerAssignment is also the first tool that supports assignment of the recently developed copy number signatures^25^, which are good predictors of clinical survival^25,26^.

In summary, SigProfilerAssignment provides a novel computational package and an accessible online interface to accurately assign known mutational signatures to an individual cancer and individual somatic mutations, thus, enabling users to ascertain the mutational processes operative in a cancer genome.

### Algorithm 1 Assigning mutational signatures to samples with SigProfilerAssignment

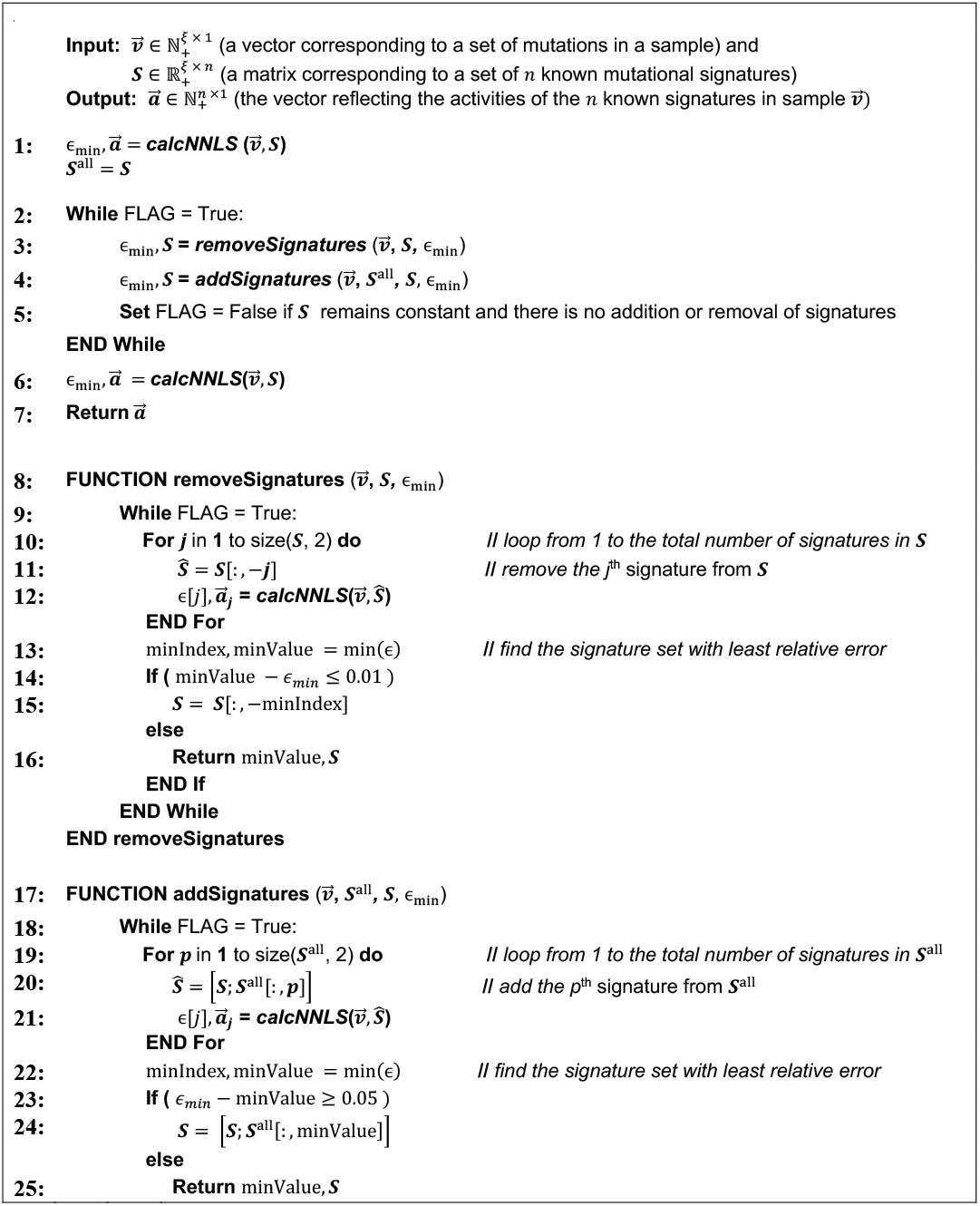
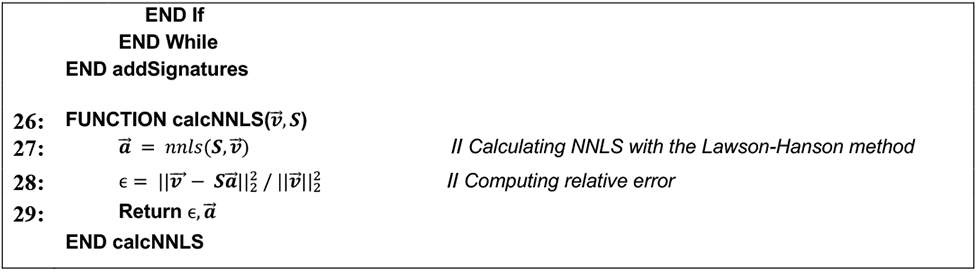

## ONLINE METHODS

### Distribution and Usage

SigProfilerAssignment is distributed as a Python and R package (https://github.com/AlexandrovLab/SigProfilerAssignment and https://github.com/AlexandrovLab/SigProfilerAssignmentR) with support for most operating systems and an extensive documentation at https://osf.io/mz79v/wiki/home/?view_only=5998aee28bd14ef6bda9189184a984bd. In addition, a user-friendly online interface is provided as part of the COSMIC Mutational Signatures website^10^ at https://cancer.sanger.ac.uk/signatures/assignment/. For compliance with EU and UK specific privacy regulations, the COSMIC website requires free registration prior to using SigProfilerAssignment. This ensures that all uploaded user data are maintained privately and purged properly.

Input data for both desktop and online versions can be provided by mutation calling and segmentation files, depending on the variant class, and is processed internally by SigProfilerMatrixGenerator^22,27^. The tool supports common formats for SBS, DBS, and ID somatic mutations, including the Variant Call Format (VCF), the Mutation Annotation Format (MAF), and simple text files. Multi-sample segmentation files obtained from ASCAT^28^, ABSOLUTE^29^, Sequenza^30^, FACETS^31^, Battenberg^28^, or PURPLE^32^ are supported for analysis of copy number signatures. In addition, SigProfilerAssignment can use standard mutational matrices, where rows correspond to mutational channels and columns to samples, extracted from the SigProfiler suite of tools^16,22,33^. Different sequencing assays (whole genome sequencing, whole exome sequencing, and targeted sequencing), species (human, mouse, and rat), genome builds (GRCh37/38, mm9/10, and rn6), and signatures (default COSMICv3.3^10^, prior COSMIC versions, and custom signature databases) are supported.

The main output of SigProfilerAssignment includes the activity of each known mutational signature for each of the supplied samples, the reconstruction of the original dataset, and the probability of each individual mutation being caused by a specific signature. The latter is not provided when the input file is a mutational vector or mutational matrix as this input format lacks information about individual somatic mutations. Signature activities correspond to the specific numbers of mutations from the original catalog caused by a particular mutational process. Considering these activities, as well as the provided set of known mutational signatures, a reconstruction of the original mutational catalog for each sample is derived. Different accuracy metrics for this reconstruction are outputted by SigProfilerAssignment, including cosine similarity, Kullback–Leibler divergence, Pearson correlation, L1 relative error, and L2 relative error.

The signature assignment results are summarized using three independent visualizations: *(i)* a bar plot depicting the activities of all mutational signatures within a sample; *(ii)* a tumor mutational burden (TMB) signature plot showing the activities per mutational signature; and *(iii)* an individual reconstruction plot per sample, which includes the mutational profiles for both the original and the reconstructed input sample, different accuracy metrics, and the mutational profiles for each of the known mutational signatures assigned to that sample. For the online version of the tool, an interactive heatmap plot, including the signatures’ activities and the samples’ reconstruction accuracies is also provided. Raw data files containing activities, reconstruction metrics, and signature probabilities for individual mutations are generated by the desktop tool and can be downloaded from the online version.

### Benchmarking of bioinformatics tools for refitting known mutational signatures

To evaluate the performance of tools for refitting known mutational signatures, we used a standard set of evaluation metrics and compared SigProfilerAssignment with another four commonly used approaches: deconstructSigs^17^, MutationalPatterns^18,19^, sigLASSO^20^, and SignatureToolsLib^5,21^. Specifically, each tool was applied to 2,700 previously simulated cancer genomes^16^, corresponding to 300 simulated tumors from nine different cancer types, including: bladder transitional cell carcinoma, esophageal adenocarcinoma, breast adenocarcinoma, lung squamous cell carcinoma, renal cell carcinoma, ovarian adenocarcinoma, osteosarcoma, cervical adenocarcinoma, and stomach adenocarcinoma. The cancer genomes of these samples were simulated using 21 different COSMIC reference signatures. To emulate a typical refitting of mutational signatures, each tool was applied by utilizing the complete set of 79 COSMICv3.3 SBS signatures. After assigning the signatures, the assignment of each signature to each sample was classified as either a *true positive* (TP), *false positive* (FP), or *false negative* (FN) result. A known signature was considered TP if at least one mutation was assigned to the signature by a particular tool and the ground truth activity of the signature was greater than zero. In contrast, a signature was classified as FP when it was assigned by a tool, but the ground truth activity was zero. Lastly, FN results were signatures with ground truth activities above zero that were not assigned any somatic mutation. These standard metrics allowed calculating the precision, sensitivity, and F_1_ score of each tool per sample, defined as:

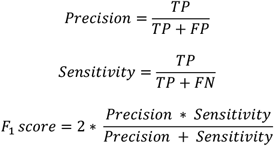

These metrics were calculated for each synthetically generated sample and, subsequently, averaged to obtain a final accuracy value for each random noise level (0%, 5%, and 10%). To benchmark the computational performance of the different bioinformatics tools, their CPU elapsed time and peak memory usage were monitored and averaged for the three noise levels.

SigProfilerAssignment v0.0.28 was run using default parameters. deconstructSigs^17^ v1.8.0 was used with default parameters as indicated in https://github.com/raerose01/deconstructSigs/. MutationalPatterns^18^ v3.0.1 was run with default parameters independently using its standard and strict modes, corresponding to the *fit_to_signatures* and *fit_to_signatures_strict* functions, respectively. The *max_delta* parameter was fixed to a default value of 0.004 for the strict mode, according to authors’ instructions at https://bioconductor.org/packages/release/bioc/vignettes/MutationalPatterns/inst/doc/Introduction_to_MutationalPatterns.html. sigLASSO^20^ v1.1 was used with default parameters (no priors) following the instructions at https://github.com/gersteinlab/siglasso; albeit avoiding the generation of plots for the comparison of the computational performance. SignatureToolsLib^5^ v2.1.2 was run with global signatures using the *Fit* function and default parameters as indicated at https://github.com/Nik-Zainal-Group/signature.tools.lib.

## DATA AVAILABILITY

All synthetic benchmarking data used in this article are available on FigShare at https://doi.org/10.6084/m9.figshare.20409430 and were originally generated as part of Ref. ^16^. They are publicly available under the Creative Commons Attribution 4.0 International license.

## CODE AVAILABILITY

SigProfilerAssignment is developed as a Python package and it is available under a permissive BSD 2-clause license at https://github.com/AlexandrovLab/SigProfilerAssignment and https://pypi.org/project/SigProfilerAssignment/. An R wrapper is also provided using the same license at https://github.com/AlexandrovLab/SigProfilerAssignmentR. SigProfilerAssignment provides support for most operating systems, including Windows, macOS, and Linux-based systems. An online version of the tool, requiring a free registration, is provided as part of the COSMIC Mutational Signatures website at https://cancer.sanger.ac.uk/signatures/assignment/.

## ACKNOWLEDGEMENTS

This work was supported by Cancer Research UK Grand Challenge Award C98/A24032, as well as US National Institute of Health grants R01ES030993-01A, R01ES032547, and R01CA269919, and a Packard Fellowship for Science and Engineering to LBA. The funders had no roles in study design, data collection and analysis, decision to publish, or preparation of the manuscript. The computational analyses reported in this manuscript have utilized the Triton Shared Computing Cluster at the San Diego Supercomputer Center of UC San Diego.

## AUTHOR CONTRIBUTIONS

The tool was conceptualized by LBA, MDG, and RV with assistance from PB and MRS. MDG and RV developed the Python and R code with assistance from MB, SMAI, and IV. IV deployed the online application as part of the COSMIC Mutational Signatures website, with assistance from MDG, RV, and MB. The code was extensively tested by MDG, RV, TY, NBN, SM, and SS including multiple rounds of refinement. MDG and RV documented SigProfilerAssignment with assistance from MB, TY, and ZJ. MDG, RV, and XW performed the benchmarking of all signature assignment tools on synthetic data, with assistance from MB and NBN. MDG and RV wrote the manuscript. LBA supervised the overall development of the code and writing of the manuscript. All authors read, provided input, and approved the final manuscript.

## COMPETING INTEREST

LBA is a compensated consultant and has equity interest in io9, LLC and Genome Insight. His spouse is an employee of Biotheranostics, Inc. LBA is an inventor of a US Patent 10,776,718 for source identification by non-negative matrix factorization, and he also declares U.S. provisional applications with serial numbers: 63/289,601; 63/269,033; 63/366,392; 63/412,835; 63/483,237; and 63/492,348. All other authors declare no competing interests.

